# Attention to Psuedo-Tone Melodies Enhances Cortical but Not Brainstem Responses in Humans

**DOI:** 10.64898/2025.12.15.694407

**Authors:** Victoria Figarola, Yunshu Li, Adam Tierney, Fred Dick, Abigail Noyce, Ross K Maddox, Barbara Shinn-Cunningham

**Author notes:** Equal Contributions (Victoria Figarola, Yunshu Li). **Conflict of interest statement:** The authors declare no competing financial interests.

## Abstract

Auditory selective attention, the ability to focus on specific sounds while ignoring competitors, enables communication in complex soundscapes. Though attention clearly modulates cortical responses to sound, whether and where this modulation occurs in subcortical structures remains disputed.

Here, we use electroencephalography to record cortical and subcortical (auditory brainstem responses, ABRs) activity during a selective attention task. Human participants attend to a 3-note melody in one pitch range presented to one ear while ignoring a competing, interleaved melody in a different pitch range played to the other ear (Laffere et al., 2020, 2021). The melodies consist of pitch-evoking pseudo-tones formed by convolving a periodic impulse train with a brief tone pip. These stimuli allow us to measure both ABRs (elicited by each individual tone pip within a pseudo-note) and cortical responses (elicited by the onsets of pseudo-notes) simultaneously.

We observed robust ABRs, but no evidence of modulation by attention. Conversely, cortical responses, measured by event related potentials (ERPs), demonstrated attentional modulation of the P1-N1 peak. We conclude that attentional modulation within the brainstem is not measurable in the well-defined peaks of the ABR, which themselves reflect processing up to the input stage to the inferior colliculus.

## 1 Introduction

Listening in noisy settings requires top-down attention to enhance neural representations of task-relevant sounds (Cherry, 1953; Bronkhorst, 2000; Best et al., 2008; Shinn-Cunningham and Best, 2008; Hill and Miller, 2010). Individuals with hearing impairments have trouble deploying top-down attention and struggle to select and maintain focus on relevant sounds amid background noise (Ruggles et al., 2012; Bharadwaj et al., 2014; Bonacci et al., 2019). Despite its clinical relevance, it is not well understood where along the auditory pathway—cortical versus subcortical—attentional modulation emerges. Here, we simultaneously probe attentional modulation of both cortical and brainstem responses to characterize attention effects throughout the auditory pathway.

Electroencephalography (EEG) responses can assay neural processing and attention effects throughout the auditory system, including attentional modulation of auditory information. Cortical event-related potentials (ERPs) evoked by a particular sound are enhanced when that sound is attended compared to when it is ignored (Hillyard et al., 1973; Choi et al., 2014; Noyce et al., 2023). EEG also can measure auditory brainstem responses (ABRs), which reflect peripheral auditory function within 10 ms following sound onset. ABR waves reveal responses in successive subcortical structures: Wave I (distal auditory nerve), Wave II (proximal auditory nerve), Wave III (cochlear nucleus), and Wave IV and Wave V (overlapping contributions from rostral brainstem structures, including the superior olivary complex, lateral lemniscus, and inferior colliculus) ((Eggermont, 2019)). ABRs thus can be used to evaluate whether attention modulates neural activity within the auditory brainstem.

While cortical attentional effects are well established, effects on sound processing at a subcortical level remain debated. Some studies propose that early auditory nuclei filter out irrelevant inputs (Lukas, 1980; Hackley et al., 1990; Maison et al., 2001). In non-human animals, inhibition of auditory responses occurs in the cochlear nucleus and inferior colliculus (cats: (Hernandez-Peon et al., 1956; Desmedt, 1962; Oatman, 1971; Oatman and Anderson, 1977; Kawase et al., 1993); ferrets: (Bajo et al., 2007; Slee and David, 2015); monkeys: (Ryan and Miller, 1977); guinea-pig: (Nieder and Nieder, 1970; Dolan and Nuttall, 1989)). However, most past studies examined task-driven modulation in the absence of distractors or competing stimuli rather than during selective attention. This distinction is important: without competing streams, observed effects may reflect global arousal, vigilance, or task engagement rather than top-down attentional selection.

In humans, claims of attentional modulation in subcortical responses are inconsistent. While some studies report effects in brainstem responses (Forte et al., 2017; Etard et al., 2019; Mankel et al., 2024; Strauss et al., 2025) or the inferior colliculus (Rinne et al., 2008; Etard et al., 2019), others find no modulation of the ABR or the auditory steady-state response (ASSR) (Connolly et al., 1989; Woldorff and Hillyard, 1991; Rinne et al., 2007; Varghese et al., 2015; Stoll et al., 2025). Critically, many of these studies rely on indirect measures with limited temporal precision (e.g., fMRI or ASSRs), making it impossible to attribute effects to specific subcortical generators whose responses overlap with each other and with cortical responses. Given these methodological limitations and inconsistent replication across studies, effects of attention on human subcortical auditory processing remains an open question.

We combined two established EEG paradigms (Laffere et al., 2020, 2021); (Polonenko and Maddox, 2019, 2021; Laffere et al., 2020, 2021) to simultaneously measure neural responses from both the brainstem and cortex as participants engaged top-down selective attention. Participants heard competing, temporally interleaved low- and high-pitched notes presented to different ears. The notes were engineered so that 1) low and high melodies excited distinct cochlear regions, 2) each note elicited multiple ABRs, and 3) note onsets evoked cortical responses. This design allowed us to test for attentional modulation across the auditory pathway; in particular, we could look for attention modulation of subcortical responses during a task demanding enough to elicit robust attentional modulation of cortex.

## 2 Materials and Methods

### 2.1 Participants

Younger adults (18-35 years old) were recruited from the Carnegie Mellon University community and were compensated for their time in either cash or partial course credit. Participants were screened to have no diagnosed hearing impairment or neurological disorder affecting auditory function, to have pure-tone audiometric thresholds within 20 dB of normal (125 Hz to 8000 Hz), and to be able to perform the experimental task (detailed under Training below).

Rather than using a power analysis to determine the number of participants, we adopted a Bayes Factor approach, collecting data until we had enough subjects to support either finding or rejecting attentional modulation of subcortical responses (see Statistical Methods, below). Based on this approach, we recruited 53 participants. Of these, 36 passed all screenings and the training task and completed the full experiment (18 men, 17 women, 1 gender not reported; age 22.13 ± 3.73 years), while 10 participants failed the hearing screening, 5 failed the training, and 2 were dismissed due to technical issues.

### 2.2 Stimulus Design

Each trial consisted of six notes, alternating between low and high pitch to create an isochronous sequence consisting of a three-note “low” melody temporally interleaved with a three-note “high” melody (**Figure 1**). Trials were concatenated together, separated by a silent interval with the duration of approximately two notes, to create experimental blocks (see below for further details). To support perceptual segregation of the two melodies within each trial and block, notes were randomly selected (with replacement) from pitches corresponding to the first three notes of an F-major scale (low) and the D-major (high) scale starting nine semitones above the F-major scale. Specifically, we used pitch frequencies of F1, G1, and A1 (43.65Hz, 49Hz, and 55Hz) in the low-pitch melody and D2, E2, and F#2 (73.42Hz, 82.41Hz, and 92.5Hz) in the higher-pitch melody. The perceptual pitch of each note was determined by the temporal repetition rate of the pip train rather than by resolved harmonics. Because notes were drawn with replacement within each three-note melody, identical pitch repetitions (e.g., A-A-F) could occur at any position within the triplet. This randomization was identical across Attend High and Attend Low conditions, so any effects of within-triplet repetition on ERP magnitude would apply equally to attended and ignored notes and thus do not bias our within-note attentional contrasts.

**Figure 1.**
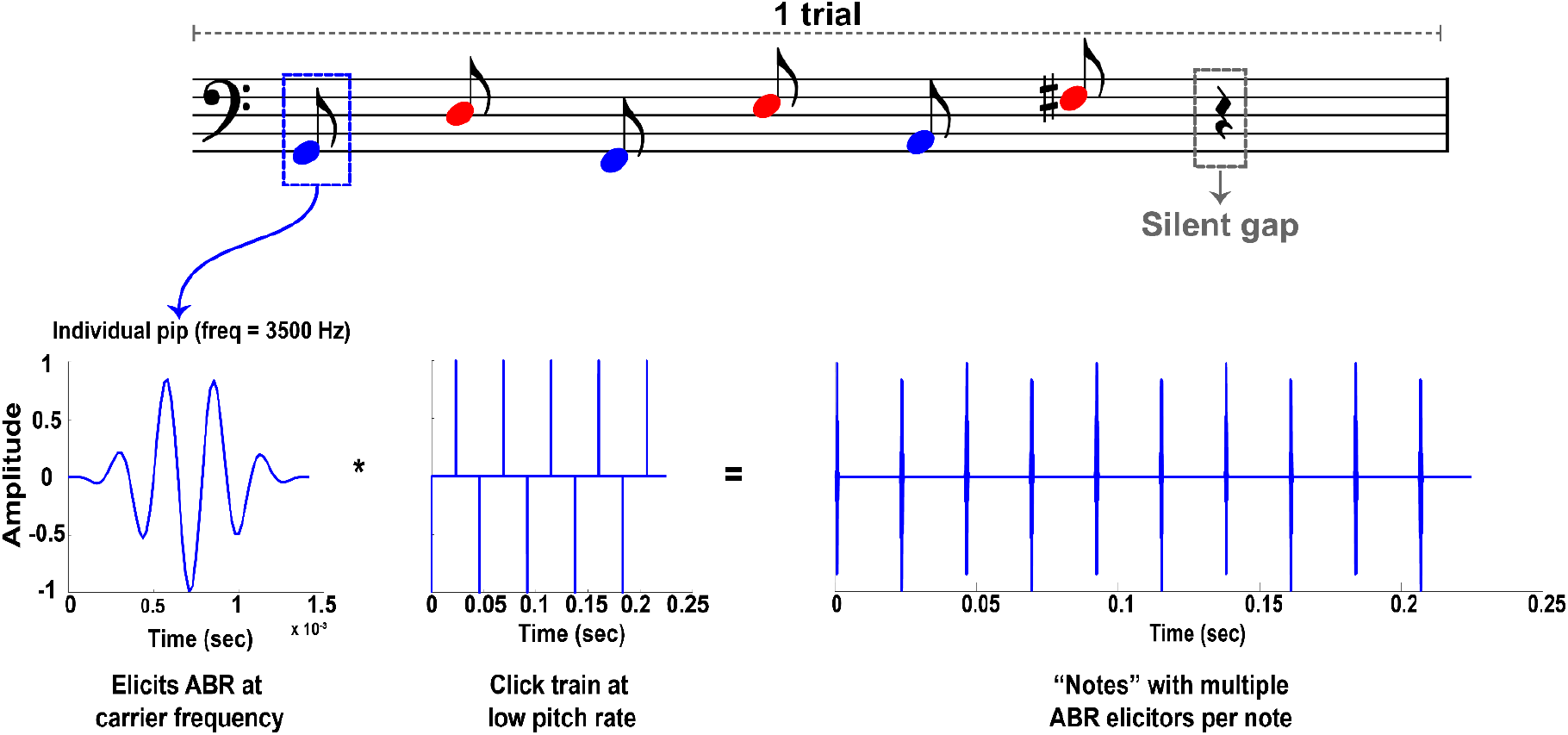
Stimulus Generation. A single trial comprises six notes, 3 high and 3 low. In this example, the tone pip repetition rate corresponding to the perceived pitch of the first note is 49 Hz. A windowed cosine tone pip (carrier frequency 3500 Hz) is convolved with a periodic impulse train (frequency 49 Hz) to generate the pitch-conveying pseudo-tone. When the note is presented to a listener, each tone pip elicits an ABR, while the pseudo-note onset elicits a single cortical onset response.

To further facilitate perceptual segregation, the low melody was always presented to one ear (left), and the high melody to the other (right). Importantly, the two streams also differed in their tone-pip carrier frequencies (CF; 3500 Hz for low notes; 4500 Hz for high notes). This choice was intentional: because ABR generators are strongly tonotopically organized, using distinct carrier frequencies helps ensure that each melody activates partially non-overlapping cochlear regions, thereby improving the signal to noise ratio of the brainstem responses and minimizing interference when extracting stream-specific ABRs. Additionally, at these high carrier frequencies, envelope periodicity (not carrier frequency) determines perceived pitch, so the CF manipulation does not meaningfully contribute to perceptual segregation.

### 2.3 Construction of Psuedo-Tone Notes

Each note within the melodies was created by convolving a narrowband tone pip with an impulse train (each impulse is a single sample set at 1) at that note’s desired pitch (fundamental frequency), creating a “pseudo-tone.” Each tone pip was a cosine of the appropriate carrier frequency multiplied by a Blackman window, with the length of the window determined by the carrier frequency (each window spanned five cycles of the tone pip’s center frequency). Within the impulse train, every other impulse was inverted to reduce stimulus artifacts from consistently positive-biased stimuli. The resulting pseudo-tone note thus comprised a series of brief tone pips with alternating polarity whose onsets were temporally spaced according to that note’s desired pitch, similar to stimuli that evoke pitch percepts using amplitude-modulated noise (Burns and Viemeister, 1981). Stimuli were generated in MATLAB with a 48 kHz sampling rate using custom code written by the authors. To facilitate reproducibility and allow readers to hear the perceptual quality of these pseudo-tone melodies, we provide example audio files (a single trial) as Supplementary Audio S5.

Each note had a duration of 330 ms and the presentation rate was one note onset every 340 ms (overall presentation rate of 2.94 Hz; 1.47 Hz within each of the low and high melodies). A silent gap of 700 ms followed each six-note trial; thus, each trial was 2.74 seconds long (2.04 seconds of interleaved notes followed by 700 ms of silence). Within each condition (Attend High & Attend Low), participants completed 16 blocks of 20 trials, for a total of 320 trials per condition. The order of conditions was counterbalanced between participants.

When a pseudo-note is presented, its onset elicits a clear cortical ERP, but each individual tone pip within a pseudo-note elicits an ABR (Bauch et al., 1980; Kileny, 1981; Hayes and Jerger, 1982) This design produces an order of magnitude more ABR responses than cortical responses, since each pseudo-note contains tens of tone pips, allowing us to obtain good estimates of ABRs in the same time needed to get good cortical ERP estimates, even though ABRs have a low signal-to-noise ratio.

### 2.4 Behavioral Task

In each block of trials (see Experiment-specific details below), listeners attended to one pitch range presented to one ear (e.g., low-pitched notes in the left ear) while ignoring the interleaved melody presented to the other ear in the other pitch range. Pilot testing showed that the low melody pitches were more challenging to hear. Given this, and the fact that we were interested in comparing neural responses to identical stimuli differing only in the focus of attention, we did not counterbalance which melody started first; instead, all trials always began with the presentation of a low note (in the left ear). Participants were instructed to identify one-back repeats of the 3-note melodies in the target pitch range and to press a button whenever they heard a melody identical to that on the immediately preceding trial (**Figure 2**).

**Figure 2.**
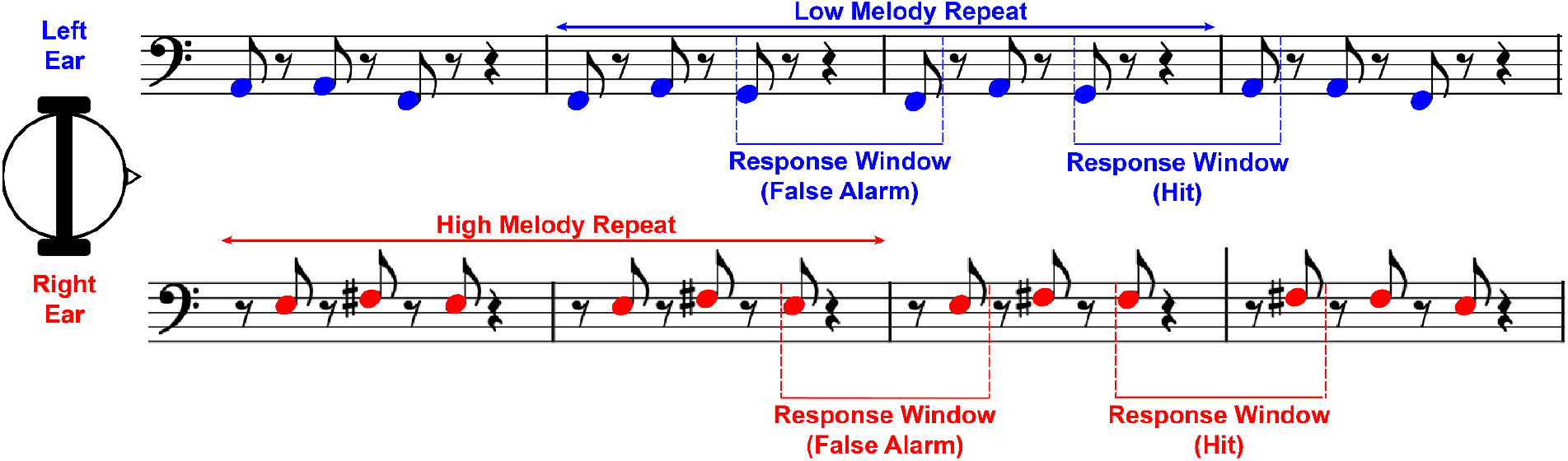
Behavioral Task. Conceptual schematic of the auditory 1-back task. *This illustration is indented as a musical schematic and is not drawn to scale with respect to exact stimulus duration or inter-stimulus intervals*. Stimuli consisted of interleaved low- and high-pitched pseudo-notes, separated by approximately one octave. Participants were asked to respond whenever a 3-note pattern within the target melody was identical to that of the preceding trial. Button presses were scored as hits when they occurred between the onset of the last note of the repeated trial and the offset of the 1st note of the next trial, and as false alarms in that time range on non-repeated trials; otherwise, they were discarded.

### 2.5 Experimental Procedure

After passing an initial screening, all participants participated in a 30-minute training paradigm (implemented on https://gorilla.sc; (Anwyl-Irvine et al., 2020)) that taught them how to selectively attend to either the high or low interleaved tone pattern. This training was necessary because perceiving the pitches of the pseudo-tones can be challenging, particularly for low-pitched notes (Plack et al., 2005), especially if individuals are not familiar with the stimuli. To enter the full study, participants had to achieve a performance of *d’* = 0.7 or higher on the training task in both high and low conditions. Participants were given up to 3 chances to repeat the training to achieve this score (see experiment-specific methods).

Participants who passed all screenings and successfully completed the training phase were invited to complete the full experiment. EEG electrodes were applied (as described below) and participants were seated in a sound-attenuated room. Stimuli were presented via ER-3A insert earphones (Etymotics) with a sound presentation level of 65 dB SPL.

All experimental control scripts were written in MATLAB Psychtoolbox (Kleiner, M., Brainard, D., Pelli, D., Ingling, A., Murray, R., & Broussard, C, 2007). The start of each experimental block began with a cue to direct the participants’ attention, consisting of an isolated pair of 3-note trials presented in the target pitch band and the appropriate ear for that pitch band (i.e., just like the trials in the upcoming block, but without the competing notes in the other ear). Throughout the block, a small cross was presented in the center of the screen to facilitate steady visual fixation, reducing ocular artifacts in the EEG.

### 2.6 EEG Data Acquisition

EEG data were recorded on a BioSemi ActiveTwo 32-channel EEG system (electrodes positioned according to the 10/20 montage; sampling rate 16,384 Hz) using Ag-AgCl active electrodes. Data were also recorded from two external reference electrodes placed on the earlobes. Offsets were maintained beneath 20 millivolts. An additional sound card channel was used to align EEG event markers (triggers) with the start of each tone pip for subsequent analyses; these event markers were recorded by the ActiveView software simultaneously with the EEG data.

### 2.7 Behavioral Analysis

Behavioral performance was quantified using listeners’ discrimination (*d’*) between repeats and non-repeats. Keypress responses were counted as *hits* when they fell between the onset of the final note of a repeated target melody and the offset of the first note in the target melody of the next trial, and as *false alarms* when they fell into the same time window following a non-repeated melody (**Figure 2**). Keypress responses falling outside of these two windows were discarded.

### 2.8 EEG Analyses

All analyses were conducted via custom MATLAB scripts using EEGLab (Delorme and Makeig, 2004).

### 2.8.1 Subcortical: Auditory Brainstem Response (ABR)

Only channel Cz was analyzed, as the dipoles for subcortical structures, particularly those generating ABR Wave V, are oriented for optimal signal at Cz (Parker, 1981; Eggermont and Schmidt, 1990; Skoe and Kraus, 2010). Data were re-referenced to the average of the earlobes. Event markers were shifted 1 ms later to account for air tube delay induced by the ER-3A earbud system. A first-order causal, Butterworth bandpass filter (100–1500 Hz) was applied to isolate the ABR components, and the data were then subsequently epoched from -5 to 15 ms relative to the onset of each individual tone pip. Consistent with standard ABR practice, no baseline correction was applied. To confirm that the absence of baseline correction did not influence the results, we repeated analyses using 1-, 3-, and 5-ms pre-stimulus baselines (Supplementary Figure S1).

To reduce overall noise and most effectively estimate the ABR signal, we adopted the ABR response-averaging method outlined by (Polonenko and Maddox, 2019). Leveraging the fact that the ABR signal is approximately 100 times weaker than noise (such that the main contribution to variance in each trial could be attributed to noise), we estimated the reliability of the ABR elicited by each individual tone pip from the variance calculated across the entire 6-note trial in which that tone pip appeared. Weights were computed and applied within each attention condition, ensuring attended and ignored trials were never mixed during weighting. Weighted epochs for each condition (Attend High, Attend Low) were then averaged separately for each ear. For completeness, we also computed ABRs using standard (non-weighted) averaging procedures. These results are in Supplementary Figure S2.

For each participant, Wave I, Wave II, Wave III, Wave V, and Wave VI amplitudes were defined by averaging the voltage within time windows that we estimated from the grand average waveforms across all participants: Wave I (1.5-2.5ms), Wave II (3-3.5ms), Wave III (3.6-4.5ms), Wave V (5–6.5 ms) and Wave VI (7-8.5ms).

#### 2.8.2 Cortical: Preprocessing

EEG data were downsampled to 500 Hz. A zero-phase, 1^st^ order Butterworth bandpass filter was applied from

0.3 (to minimize slow drift) to 30 Hz (to minimize line noise and myogenic artifact). Next, the data were decomposed using independent component analysis (ICA); components corresponding to eye blinks, eye saccades, and motor-related artifacts were identified (by visually inspecting the topographies and time courses, with reference to automated ICA labeling (Makeig et al., 1995) and removed.

#### 2.8.3 Cortical: Time Domain Analysis

The data were epoched around the start of each trial (see specific research methods below) and averaged across 14 fronto-central channels (channels Fp1, AF3, F3, FC1, FC5, C3, CP1, C4, FC6, FC2, F4, AF4, Fz, and Cz after (Laffere et al., 2020)). Each epoch was baseline corrected by subtracting the average across a window from -100 to 0 ms relative to stimulus onset. Trials were then epoched from -0.10 to 2.74 seconds relative to the onset of that trial’s first note.

The N1 component of an ERP is a well-established marker of early auditory cortical processing and is known to be modulated by attention (Hillyard et al., 1973; Steven J. Luck, 2000; Choi et al., 2013). In particular, we focus on how attention modulates the P1-N1 peak-to-peak amplitude, capturing the combined influence of sensory encoding (P1) (Fogarty et al., 2020) and attentional enhancement or suppression (N1) (Näätänen and Picton, 1987; Choi et al., 2014).

For the note-based ERP analysis, data were epoched from 0 to 340 ms relative to each note onset. Because our goal was to isolate attentional modulation within acoustically identical stimuli presented to the same ear, all attentional contrasts were computed *within* note type (attended vs. ignored high notes; attended vs ignored low notes) and never between low and high streams.

The three high-note onsets in each trial were averaged together. For the low notes, the first note in each trial, always a low note, was excluded from this averaging as in the absence of an ongoing distractor, its ERP amplitude is minimally influenced by attention (Choi et al., 2014). Thus, low-note ERPs reflect the second and third low onsets only.

All trials, including repeat and trials later containing a false alarm, were retained for ERP analysis in the main text. Because participants cannot know a trial is a repeat until after the sixth note, and because the behavioral response occurs only after the final note, ERP time windows of interest precede any motor activity. To confirm that including repeat trials did not influence the observed attentional effects, we performed a supplementary analysis in which repeat trials were excluded prior to averaging ERPs (Supplementary Figure S3).

Because the two melodies followed a predictable, alternating pattern, we examined slow anticipatory activity resembling a contingent negative variation (CNV; (Wöstmann et al., 2015)), specifically whether this activity differed across attention conditions or predicted behavioral performance. CNV-like activity was quantified as the mean amplitude in a -300 to 0 ms window before the note onset for each condition. Results of this supplementary CNV analysis are reported in Supplementary Figure 4.

After exploratory analyses, we found that we were consistently unable to identify P1-N1 peaks reliably in the ERPs to low note onsets. As such, all the analysis described below in peak picking apply solely to ERPs following high note onsets.

P1 and N1 peak values were identified by defining a time window that we estimated from the grand average waveforms across all participants (P1: 30 - 110 ms; N1: 80 - 150 ms). The maximum (P1) and minimum (N1) value in that window was identified, and the average ERP voltage in a 20 ms window (for P1) and 40 ms window (for N1) centered on the respective peaks was computed. For some participants where the standard automatic peak peaking failed due to irregular ERP morphologies, the P1 and N1 time windows were manually adjusted slightly to account for differences in ERP morphology. For finding the P1 peak, one participant’s window was adjusted to 40 - 110 ms; for finding the N1 peak, these windows were 80 - 160 ms for one participant and 80 - 200 ms for three participants. In addition to these altered windows, 4 participants were excluded entirely from this analysis due to a lack of clear P1 and N1 peaks. Excluding these four participants did not change any outcomes of the ERP analyses, and so they were retained in the dataset for completeness.

### 2.9 Statistical Analyses

#### 2.9.1 Subcortical: Auditory Brainstem Response (ABR)

Three statistical analyses were conducted. First, permutation t-tests (10,000 permutations, two-sided) were used to analyze attention effects on peak amplitudes within the time windows for each wave listed above (Maris and Oostenveld, 2007; Krol, 2024; Gerber, 2025)). Second, a cluster-based permutation test (significance threshold = 0.01; 10,000 permutations, two-sided) was applied across the entire ABR waveform for each ear separately to identify clusters where responses differed significantly when the evoking pulse was attended vs. when it was ignored (Maris and Oostenveld, 2007). Clusters were defined as continuous time points showing significant differences at an uncorrected threshold of p < 0.05. For each permutation (n = 10,000), the maximum cluster-level t-statistic was included in the null distribution. Lastly, Bayes factor analysis (Dienes, 2014) was conducted on peak amplitudes in each time window using the “BayesFactor” package in R (Morey, R.D., Rouder, J.N., Jamil, T., 2015) to quantify the strength of the evidence against or in favor of the null hypothesis (Null Hypothesis: attention does not modulate ABR components; (Kass and Raftery, 1995).

#### 2.9.2 Cortical: Time-Domain

A paired t-test directly tested attention effects on P1-N1 amplitudes of ERPs elicited by high notes (recall that for the low notes, the P1 and N1 ERP components could not be resolved sufficiently to compute this effect).

We wanted to test for any possible difference between the Attend High and Attend Low waveforms. We computed the Attend High-Attend Low difference wave and found its mean squared error (MSE). Within each individual participant, we then compared that observed MSE to a null distribution generated from 10,000 permutations of randomly reshuffled trial labels. The count of participants whose MSE was significantly higher than the null distribution (p < 0.05 uncorrected) was compared to the Bernoulli distribution generated from p = 0.05.

## 3 Results

### 3.1 Behavioral Performance

Participants’ performance (*d’*) was significantly above chance in both conditions (Wilcoxon signed-rank; z = 4.99, p=5.86× 10^-7^; **Figure 3**). Further, performance was significantly better in the Attend High condition than the Attend Low condition (Attend High median *d’*: 3.675, Attend Low median *d’*: 1.614).

**Figure 3.**
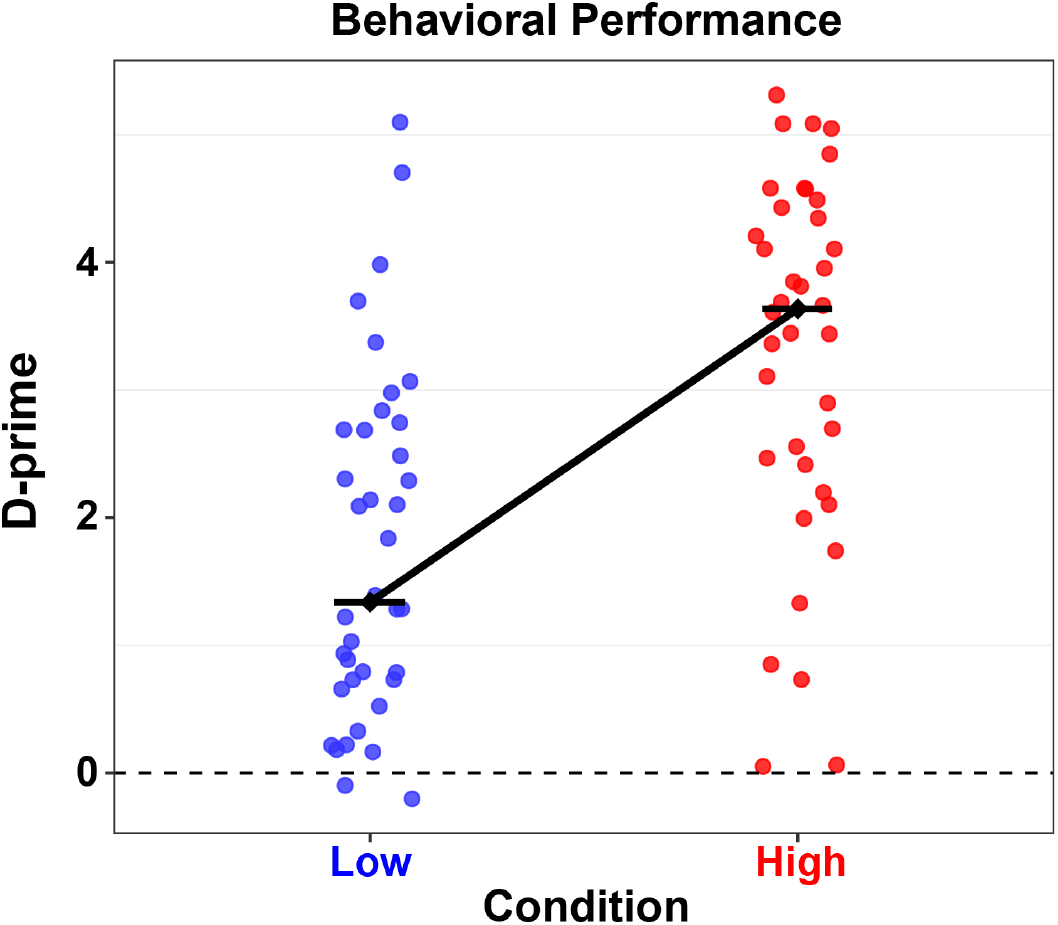
Behavioral Results. Behavioral performance is significantly higher when participants are attending to the high melody (red) than when they are attending to the low melody (blue; *p* < 0.001). The black dot and connecting line depict the median *d’* for each Attended condition.

### 3.2 No effect of attention on the ABR

We tested whether attention modulates subcortical responses by comparing ABRs elicited by acoustically identical attended and ignored stimuli within the same ear. **Figure 4** depicts the ABRs for both ears. Permutation t-tests on Wave I, Wave II, Wave III, Wave V, and Wave VI peak amplitudes revealed no significant differences between attended and ignored stimuli (all *p* > 0.13 uncorrected). A cluster permutation test (Maris and Oostenveld, 2007) across the entire ABR epoch did not identify any time points at which Attend High and Attend Low ABRs significantly differed (all p > 0.09). Bayes Factor estimates for the effect of attention on the peaks provided moderate evidence in favor for the null hypothesis (i.e., no effect of attention) for most conditions (bolded values in **Table I**; all BF < 0.274). Conditions reported in plain text correspond to inconclusive evidence (0.33 < BF < 3). Finally, there were no significant correlations between task performance and the effect of attention on ABR amplitude across all waves (Spearman’s correlation; all p values > 0.14).

**Figure 4.**
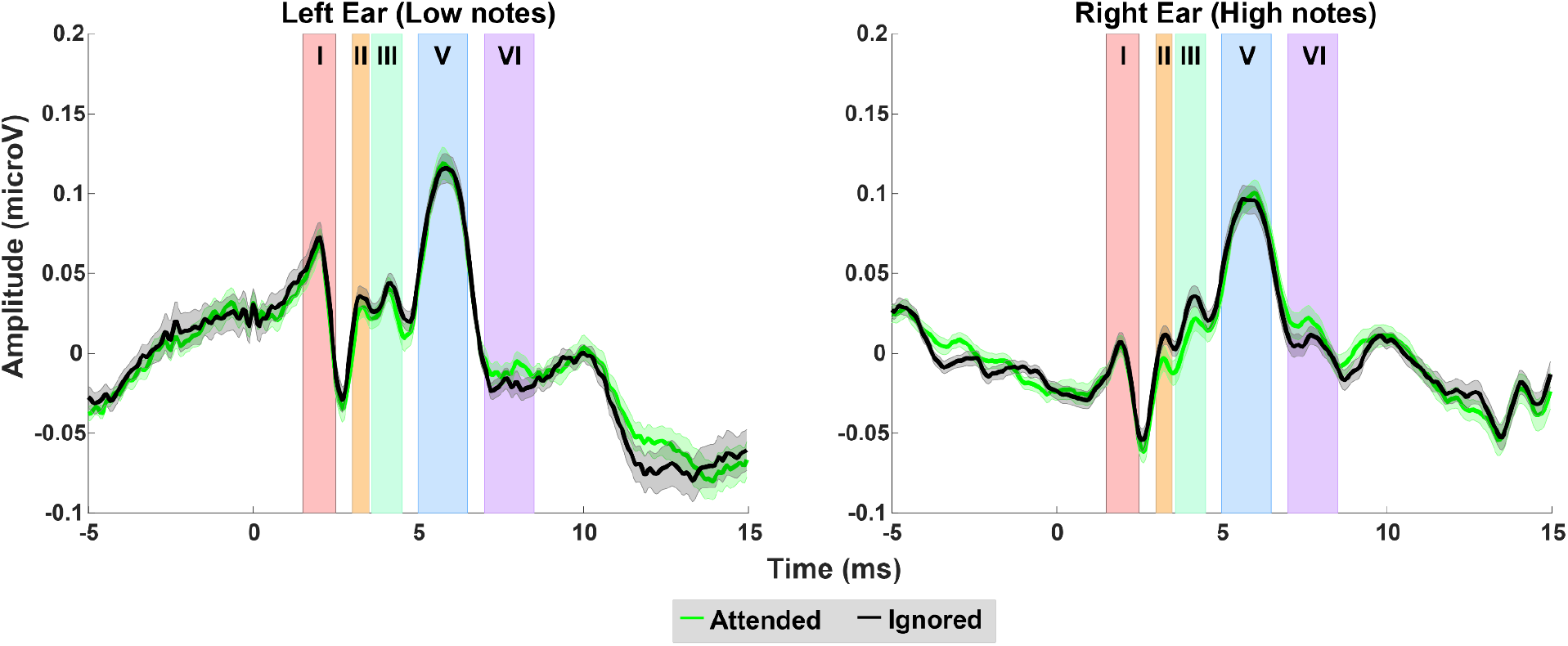
Experiment ABRs. ABRs when participants are attending to (neon green) versus ignoring (black) a given stimulus. Shading indicates time regions corresponding to Waves I-VI. Error ribbons are standard error across participants. Note that low melodies were always presented to the left ear and high melodies were always presented to the right ear. There are no significant effects of attention in any time window (all p > 0.13, all BF < 0.463).

**Table I.** ABR Bayes Factor Results. Bold refers to BFs that provide evidence in favor of the null hypothesis (BF < ⅓), and plain text refers to BFs that are inconclusive.

| **Table I.** ABR Bayes Factor Results. Bold refers to BFs that provide evidence in favor of the null hypothesis ($BF < \frac{1}{3}$), and plain text refers to BFs that are inconclusive. | | | | | |
| --- | --- | --- | --- | --- | --- |
|  | Wave I | Wave II | Wave III | Wave V | Wave VI |
| Left ear (low notes) | **0.234** | 0.387 | **0.274** | **0.141** | 0.463 |
| Right Ear (high notes) | **0.172** | 0.577 | 0.753 | **0.138** | **0.271** |

### 3.3 Cortical ERP Results

To test whether attention modulated cortical responses in across the entirety of the waveform when Attending High versus Attending Low, a bootstrapping permutation test (10,000 permutations, one-sided) was conducted on the continuous time points in the entire epoch across all individuals for each participant (refer to Experiment Methods). This test on the entire epoch revealed significant differences between attending high and attending low (z = 19.27, p<10^-6^).

The first (low) note elicits a robust P1-N1 complex regardless of attention condition (leftmost responses in the panels of **Figure 5**; consistent with prior results (Choi et al., 2014)). Subsequent high-note onsets (dashed red vertical lines in the figure panel) elicit larger peak-to-peak P1-N1 responses in the Attended High condition than the Attended Low condition. The ERP complexes elicited by subsequent low-note onsets are not as clearly structured but also appear to be modulated by attention.

**Figure 5.**
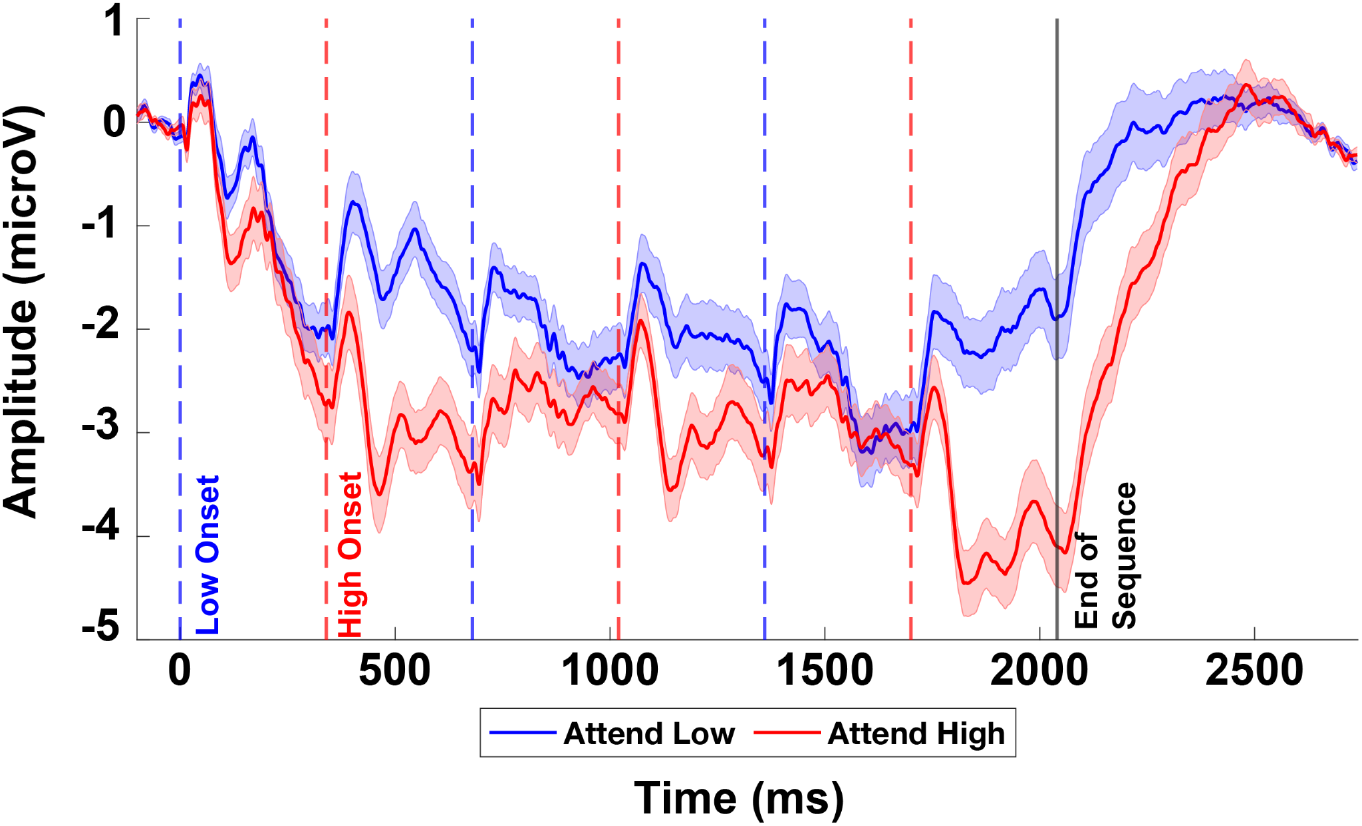
Cortical Responses. Mean EEG traces across the entire trial epoch in the Attend Low condition (blue traces) and Attend High condition (red traces). The onset of each individual note is denoted by a dashed vertical line in blue (low notes) or red (high notes). The error ribbon shows standard error of the mean within participants.

To more precisely estimate the effect of attention on cortical ERPs, we collapsed the ERP traces across the onsets to all 3 high notes, and to the second and third low notes (disregarding the first note of the trial, which was always low; **Figure 6A**) and computed the P1-N1 peak-to-peak amplitude. We were unable to identify these peaks reliably in the ERPs to low note onsets (left panel in **Figure 6A**; see Methods) so were unable to quantify attentional modulation of low note ERPs. Additionally, 4 participants’ high note ERPs were excluded from this analysis due to ERP morphologies that did not allow for a reasonable determination of their N1 and P1 peak values (refer to Methods). A total of 32 participants were included in the analysis below.

**Figure 6.**
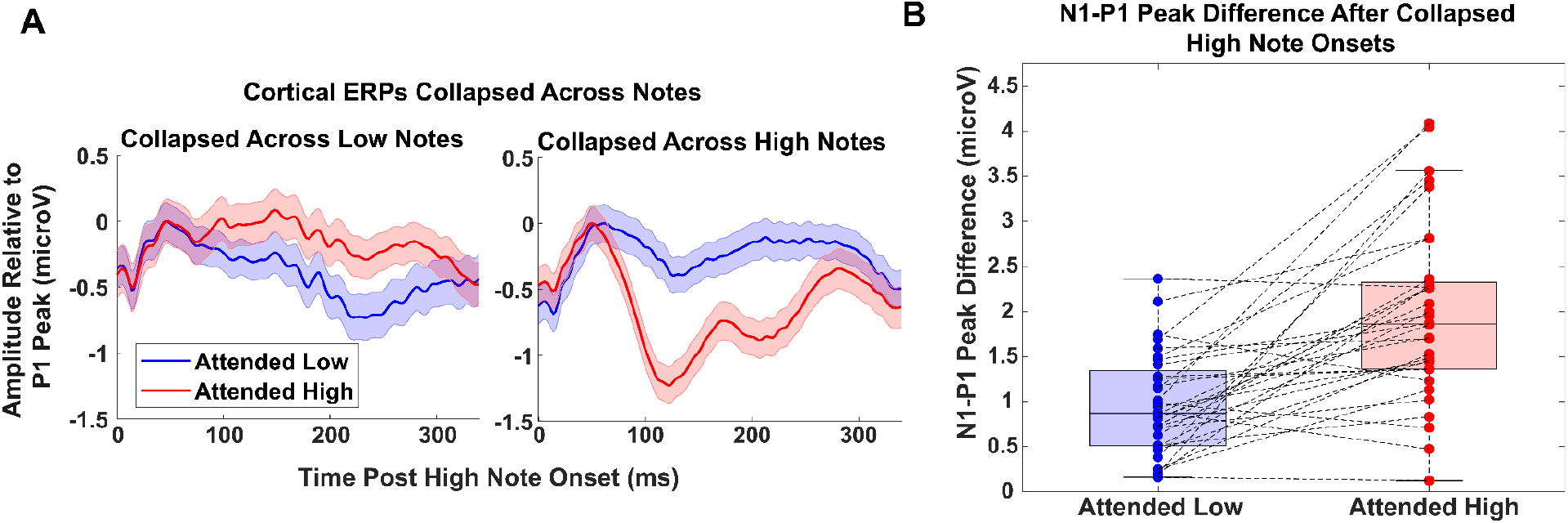
Experiment ERPs. **A:** Grand-average ERPs collapsed across the second and third low notes (left panel) and all high notes (right panel), for 32 participants with identifiable high-note P1 and N1 peaks. Low-note ERPs do not exhibit a reliable P1-N1 morphology, consistent with the reduced perceptual salience of the low-pitch psuedo-tones and the limited number of usable low-note onsets per trial. For visualization, waveforms were vertically shifted so that all P1 peaks align at 0 microV. This facilitates comparison of overall morphology but necessarily attenuates the apparent N1 amplitude due to cross-participant latency variability and peak misalignment in the grand average. Thus, the N1 in panel A reflects a latency-smeared average, not the true subject-level peak amplitude. **B:** Subject-level P1-N1 peak-to-peak amplitudes for high-note ERPs, computed from unshifted individual waveforms. Peaks were defined using windowed mean amplitudes (20 ms around P1; 40 ms around N1). These subject-specific peak estimates revealed significantly larger P1-N1 amplitudes when high notes were attended than when they were ignored (t(31) = 5.22, p = 4.81 x 10^-6^). Thus, this panel reflects the true magnitude of attentional modulation, uncontaminated by the alignment-related jitter that smooths the grand-average waveforms in panel A.

While we operationalized this effect using the P1-N1 window, it is important to note that **Figure 6A** shows a broader attention-related negativity spanning N1, P2, and N1, consistent with prior reports that attentional enhancement can extend across multiple stages of auditory cortical processing (Hansen and Hillyard, 1980). This broader negativity likely reflects the overlap of attention-related components, producing a more smeared-out negativity that was especially pronounced in the Attended High condition, in line with the behavioral performance data.

For the high-note onset responses (**Figure 6B**), a paired t-test found a significant reduction in P1-N1 amplitude in the Attended condition compared to the ignored condition post high note onset (t(31) = 5.22, p = 4.81 × 10^-6^, paired t-test). To confirm that this effect was not driven by repeat trials, we repeated the ERP analysis after excluding all repeat trials, and a significant effect of attention on the high-note onset P1-N1 remained robust t(31) = 5.693, p =2.948 x 10^-6^;paired t-test; Supplementary Figure S3).

We also explored whether the difference in P1-N1 amplitude between the attended and ignored conditions was associated with individual differences in task performance. However, this amplitude difference was not significantly correlated with performance in either task condition (Attended High Spearman’s rho = 0.1869, p_adj_= 0.6120; Attended Low Spearman’s rho = 0.2243, p_adj_= 0.4324, Bonferroni-corrected).

## 4 Discussion

We examined how attention modulates auditory processing at cortical and subcortical levels using EEG, focusing on auditory brainstem responses (ABRs) and cortical event-related potentials (ERPs). Participants selectively attended to one of two dichotically presented, interleaved three-note melodies. A critical strength of our current design is that we observed robust attentional modulation at the cortical level, confirming that attention was successfully engaged—despite the absence of subcortical effects.

### Attentional Modulation of Evoked Cortical Responses

Modulation of early cortical ERP components, especially the N1, is a well-established marker of auditory attention (Hillyard et al., 1973, 1998; Picton and Hillyard, 1974; Hansen and Hillyard, 1980; Scherg et al., 1989; Woldorff et al., 1993; Kerlin et al., 2010; Picton, 2010; Ross et al., 2010; Choi et al., 2013, 2014; Deng et al., 2019, 2020; Bonacci et al., 2020). In the present experiment, high-note onsets evoked robust P1-N1 responses, and these components were significantly enhanced when attended compared to ignored. This attentional modulation aligns with prior work showing enhanced cortical tracking of behaviorally relevant auditory objects.

On the other hand, low-note onsets elicited weaker and less clearly defined ERPs. Several stimulus- and task-relevant factors could likely contribute to this. First, the low-pitch pseudo-tones had reduced perceptual salience since their repetition rates approached the lower limit at which cortical onset synchronization is reliably elicited. Second, because the first low note was excluded and only two low-note onsets were used per trial, low-note responses were averaged over fewer samples, lowering signal-to-noise ratios. Thirdly, behavioral performance was consistently lower in the low-stream condition, suggesting that participants may have intermittently disengaged from the task, which could have further reduced the amplitude of the evoked responses. Lastly, the low stream was always presented to the same ear, raising the possibility of subtle ear-specific differences. However, any such ear-related differences cannot confound our attentional effects because all attentional contrasts were made within the same ear and within the same note type, directly comparing attended and ignored versions of the same acoustic stimulus. Additionally, this fixed mapping of each stream to a single ear is a design advantage rather than a limitation, as it stabilizes perceptual grouping of the two melodies and supports more reliable deployment of selective attention.

The predictable alternation of low and high notes also introduces the possibility of anticipatory activity. A slow negative drift resembling a contingent negative variation (CNV; (Wöstmann et al., 2015)) was present in the trial-locked average. However, additional analyses demonstrated that CNV magnitude did not significantly differ across attention conditions and did not predict behavioral performance. Thus, while anticipatory activity may shape the overall morphology of the grand average, it cannot account for the selective enhancement observed in the high-note P1-N1 responses and for the reduced low-note ERPs.

Overall, these results indicate that attentional modulation of cortical onset responses is robust for salient auditory events but is more difficult to detect for less perceptually distinct stimuli. The asymmetry between high and low notes then suggests stimulus-driven neural encoding constraints rather than a failure of attention to modulate the low stream.

### The Role of Attention in Subcortical Auditory Processing

Despite using stimuli designed to isolate brainstem activity, we found no evidence that selective attention modulates auditory brainstem responses (ABRs), including Wave V. This finding adds to a body of mixed evidence regarding subcortical attentional modulation. To address this question, our study was designed to give us the best possible chance to observe and isolate attentional modulation at each stage of processing within the brainstem, and to simultaneously measure attentional effects stemming from cortex. Specifically, by extracting clean ABRs to individual pulses, we were able to isolate and identify not only if, but where in the brainstem selective attention caused measurable effects. To achieve this, we used competing streams made up of notes whose peripheral responses could be isolated, so that overlap in the responses was negligible:

1. **Non-overlapping event times** ensured temporal separability of neural responses to low and high notes.
2. **Presentation to different ears** guaranteed that peripheral responses to low and high notes did not interact until the level of binaural convergence.
3. **Different carrier frequencies** targeted distinct cochlear regions and subsequent tonotopic channels, reducing overlap in neural responses. Importantly, these carrier-frequency differences were introduced for the sole purpose of maximizing physiological separability of the subcortical responses, not to create distinct perceptual objects. That is, they improved neural isolation of the ABR components rather than serving as cues that would meaningfully strengthen perceptual segregation.

Further, our paradigm gave listeners multiple predictable and distinct features that they could use to focus attention on the desired stream:

1. **Predictable timing** of the isochronous streams, with the low stream always starting first, allowed listeners the possibility of focusing attention in time.
2. **Presentation to different ears** further supported perceptual segregation and allowed listeners the opportunity to focus selective spatial attention.
3. **Distinct pitch ranges** for the low and high streams promoted their perceptual segregation, further supporting selective attention.

Even with these features, we found no attentional modulation of ABRs in any waves, including Wave V and Wave VI responses. Wave VI responses are typically not observed in studies using standard filtering (lowpass 1-3kHz, high pass 30-100Hz; (Picton and Hillyard, 1974; Eggermont and Schmidt, 1990; Burkard et al., 2007; Skoe and Kraus, 2010). To prevent time aliasing, we employed a narrowband filter, which allowed a Wave VI response to be isolated (Polonenko and Maddox 2022, Backer et al. 2019). Yet even this late Wave VI response showed no evidence of attentional modulation.

The absence of attentional effects suggests that top-down modulation operates above the brainstem. Our finding is at odds with some past studies; however, this may be due to differences in the paradigms and measures employed. Reports of modulation of brainstem responses from past studies using single, rather than competing sound streams could be the result of changes in arousal or task engagement (Forte et al., 2017). Past human studies reporting selective attention effects in brainstem typically used methods that cannot isolate brainstem responses in time (e.g., fMRI or FFR measures; (Rinne et al., 2008; Etard et al., 2019)), raising the possibility that slow-acting cortical responses affected these measures. However, it remains possible that subtle subcortical effects exist but are masked by neural noise.

### Behavioral Performance and the Influence of Presentation Rate and Pitch

We observed better performance in the Attend High condition. Part of this difference reflects the fact that our low notes elicited weak pitch percepts, making it difficult to track the low-note melodies. The pitch was conveyed only by temporal envelope repetition in high-frequency channels, with no low-frequency temporal fine structure cues (Oxenham, 2012). Because the repetition rate of our low notes approached the lower bound for melodic pitch perception (∼30 Hz; (Pressnitzer et al., 2001)), they conveyed a weak pitch. However, in addition, a past study observed better performance for high than low interleaved melodies, even when pure tones, rather than our tone-pip pseudo-tones, were presented (Laffere et al., 2020). Thus, it may also be generally easier to focus attention on higher-frequency melodies in this kind of interleaved-melody paradigm.

In addition to pitch limitations, the temporal structure of the stimulus also influenced performance. Specifically, interleaved melodies presented at a rapid rate can be perceived as one combined melody rather than as distinct, perceptually segregated melodies (van Noorden, 1975). This failure of segregation can occur even if the notes are presented to opposite ears, especially if pitch cues for segregation are weak (Bregman, 1990) In our paradigm, although the presentation rate was slower, the combination of weaker pitch cues in the low stream and the close temporal interleaving of the two melodies likely increased the difficulty of selectively tracking the low stream. On the other hand, high notes were more perceptually salient and more easily segregated from the distractor stream.

## Conclusion

Overall, our findings suggest that top-down attention does not significantly influence subcortical auditory processing but exerts a strong modulatory effect at the cortical level. While this suggests that attentional filtering occurs primarily after early subcortical processing stages, it may be functionally adaptive for early subcortical stages to remain unaffected by attention. Maintaining veridical representations at these levels could facilitate bottom-up redirection of attention (Näätänen, 1990) or preserve access to unattended sounds for implicit processing (Dehaene et al., 2006). These findings are consistent with late selection theories of attention, which propose that initial sensory encoding proceeds independently of attentional goals, with top-down modulation occurring at later stages (Cherry, 1953; Broadbent, 1958; Deutsch, J.A., & Deutsch, D., 1963). Future research combining EEG with techniques like MEG or fMRI could help dissociate the spatial and temporal dynamics of attentional effects across auditory hierarchies, particularly in response to naturalistic speech.

## Acknowledgements

Research reported in this publication was supported by the National Institute On Deafness And Other Communication Disorders of the National Institutes of Health under Award Number T32DC011499, the National Institute of Health Undergraduate Training Grant in Computational Neuroscience under Award Number T90DA060116. The content is solely the responsibility of the authors and does not necessarily represent the official views of the National Institutes of Health. Research reported in this publication was also supported by the Office of Naval Research (N00014-19-12332, N00014-20-12709, and N00014-23-12065), and Royal Society International Exchange Program (IES\R2\192188). We would also like to thank Audra Irvine and Andrew Levitsky for their help in data collection, and Jinhee Kim for her help on creating the musical staffs.

