## Supplemental for "Attention to Psuedo-Tone Melodies Enhances Cortical but Not Brainstem Responses in Humans"

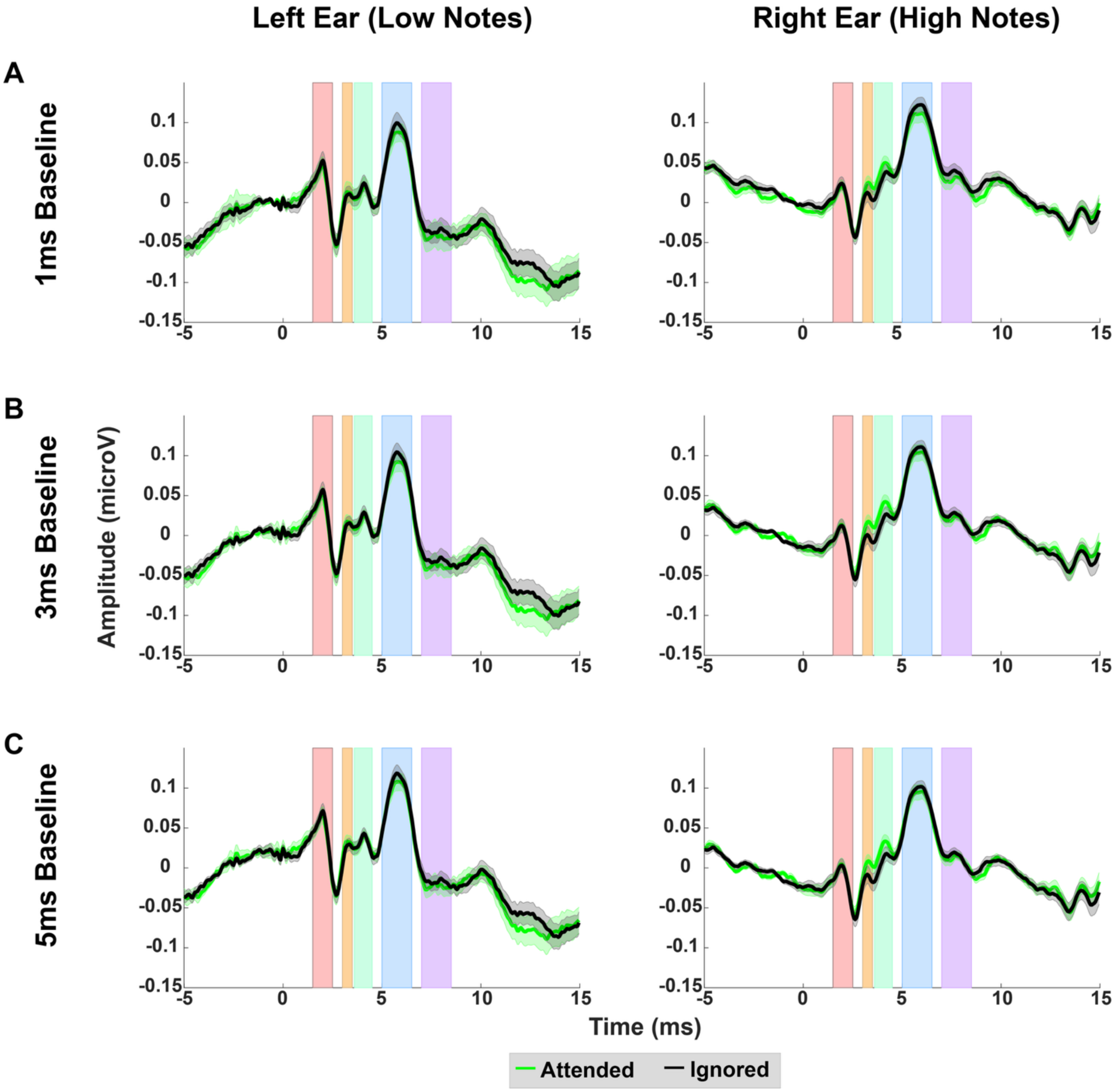


**Figure S1. ABR baseline-variant analyses.** Weighted ABRs when participants are attending to (neon green) versus ignoring (black) a given stimulus when baseline-corrected at (A) 1 ms, (B) 3 ms, and (C) 5 ms. Shading indicates time regions corresponding to Waves I-VI. Error ribbons are standard error across participants. There were no significant effects of attention in any time window or peaks across all baseline correction analysis (all $p>0.0849$).

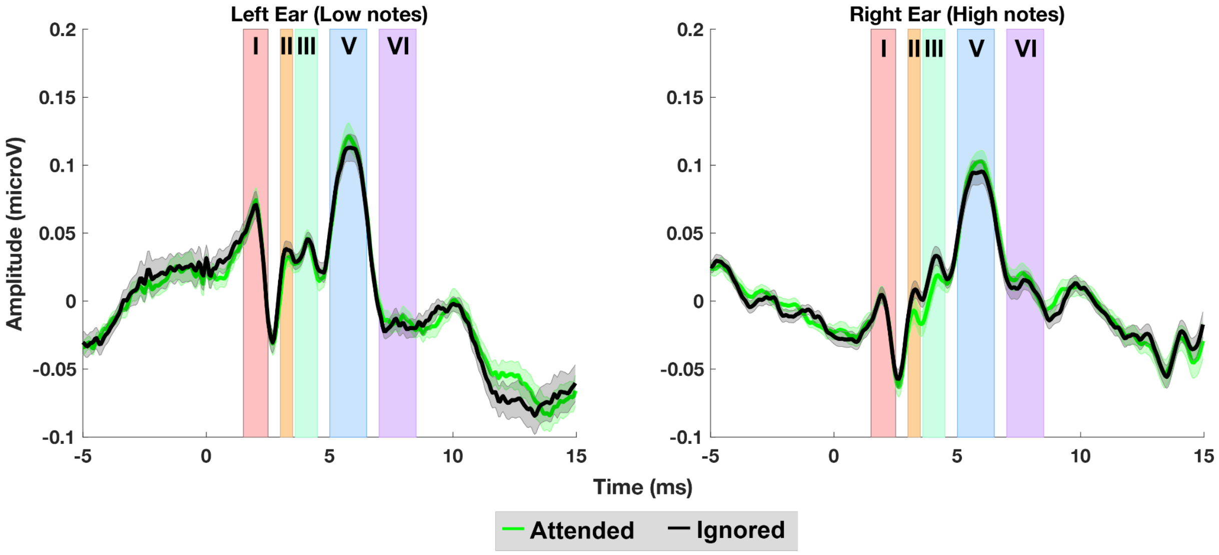


**Figure S2. Non-weighted ABRs.** Non-weighted ABRs when participants are attending to (neon green) versus ignoring (black) a given stimulus. Shading indicates time regions corresponding to Waves I-VI. Error ribbons are standard error across participants. Note that low melodies were always presented to the left ear and high melodies were always presented to the right ear. There were no significant effects of attention in any time window (all $p > 0.6769$).

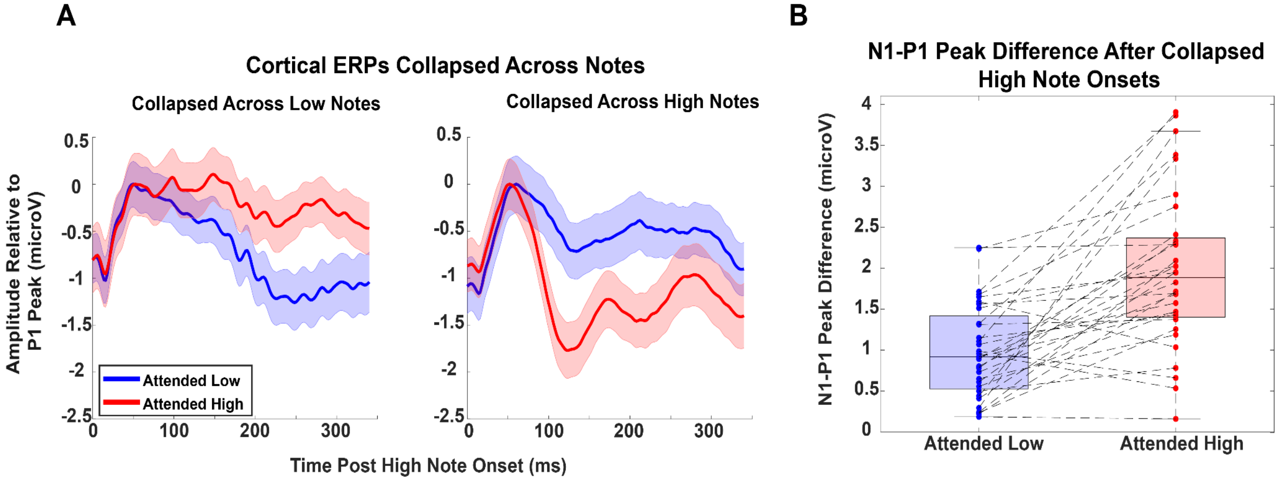


**Figure S3. Cortical ERPs and P1-N1 peak differences after exclusion of repeat trials. A:** Grand-average cortical ERPs collapsed across low-note onsets (left panel) and high-note onsets (right panel), shown separately for attended low (blue) and attended high (red) conditions. ERPs were computed after excluding all repeat trials and averaging across participants. The error ribbon shows standard error of the mean within participants. For visualization, ERP waveforms were vertically shifted so that P1 peaks align at 0 microV. **B:** Subject-level P1-N1 peak-to-peak amplitudes for ERPs elicited by high-note onsets computed from unshifted individual waveforms using windowed mean amplitudes (20 ms around P1; 40 ms around N1). These subject-specific peak estimates revealed significantly larger P1-N1 amplitudes when high notes were attended than when they were ignored ($t(31) = 5.69$, $p = 2.95 \times 10^{-6}$).

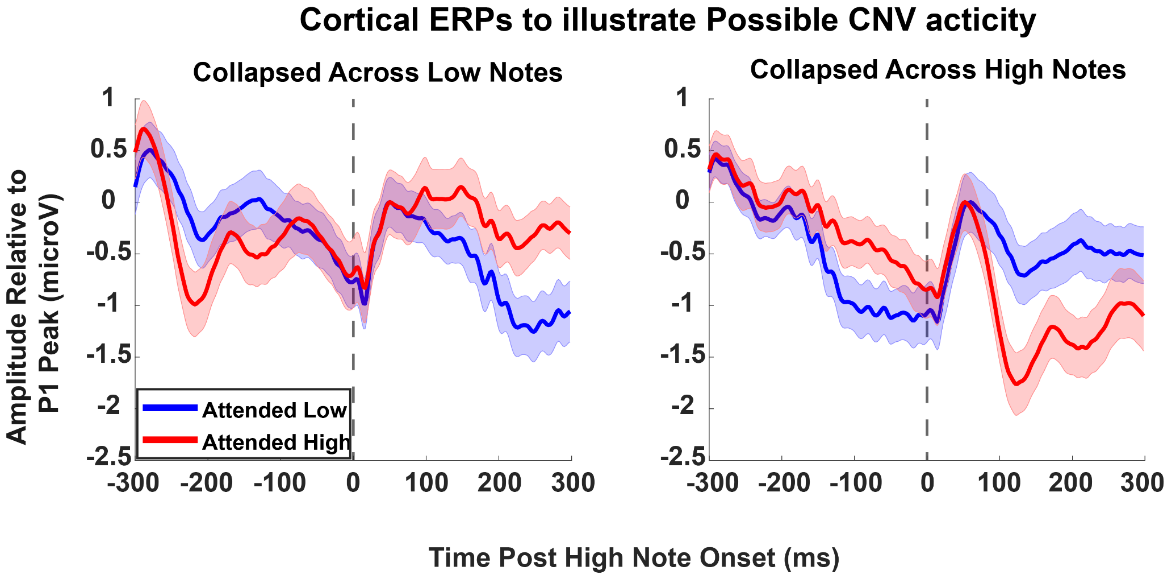


**Figure S4. CNV-like anticipatory activity does not differ across attention conditions.** Grand-average cortical ERPs time-locked to low-note onsets (panel left) and high-note onsets (panel right), shown separately for Attended Low (blue) and Attended High (red). The error ribbon shows standard error of the mean within participants. For visualization, ERP waveforms were vertically shifted so that P1 peaks align at 0 microV. CNV-like activity was quantified as the mean amplitude in the -300 to 0 ms window relative to note onset for each participant and condition. There were no significant differences in CNV amplitude between Attended Low and Attended High collapsed across the Low Notes ($t(31) = 1.953$, $p=0.1797$; paired t-test, Bonferroni-corrected) and across the High Notes ($t(31) =1.054$, $p=0.8997$; paired t-test, Bonferroni-corrected). Additionally, there were no significant correlations between CNV amplitudes and behavioral performance (all $p>0.65$; Spearman correlations). Together, these analyses indicate that while predictable stimulus timing elicits slow anticipatory activity, this CNV-like signal does not account for the attentional modulation observed in cortical onset ERPs.
